# The effects of background noise on a biophysical model of olfactory bulb mitral cells

**DOI:** 10.1101/2022.06.11.495647

**Authors:** Michelle Craft, Cheng Ly

**Author notes:** Contributing authors.

## Abstract

The spiking activity of mitral cells **(MC)** in the olfactory bulb is a key attribute in olfactory sensory information processing to downstream cortical areas. A more detailed understanding of the modulation of MC spike statistics could shed light on mechanistic studies of olfactory bulb circuits, and olfactory coding. We study the spike response of a recently developed single-compartment biophysical MC model containing 7 known ionic currents and calcium dynamics subject to constant current input with background white noise. We observe rich spiking dynamics even with constant current input, including multimodal peaks in the interspike interval distribution **(ISI)**. Although weak to moderate background noise for a fixed current input does not change the firing rate much, the spike dynamics can change dramatically, exhibiting non-monotonic spike variability not commonly observed in standard neuron models. We explain these dynamics with a phenomenological model of the ISI probability density function. Our study clarifies some of the complexities of MC spiking dynamics.

## 1 Introduction

The spiking statistics and dynamics of neural models are crucial in theoretical investigations for how animals accurately and efficiently code sensory signals. This fact has sparked numerous theoretical tools to quantify spiking behavior (Ermentrout and Terman, 2010; Gerstner and Kistler, 2002). These mathematical tools have led to great advances in our understanding of sensory neural networks, but often require making simplifying assumptions (e.g., low dimensional state variables, stable periodic dynamics, very weak or very strong noise). These limiting assumptions are easily violated when models incorporate biological realism, in which case, non-conventional or systems specific approaches must be used to better understand model behavior.

Mitral cells (**MC**) are excitatory cells in the olfactory bulb that are responsible for relaying odor information to downstream cortical regions. One of the most realistic biophysical olfactory bulb MC models was developed by the Cleland lab (Li and Cleland, 2013, 2017). Their high-dimensional multicompartment model captured known physiology of mammalian MC cells reported in experiments by several labs (Chen and Shepherd, 1997; Desmaisons et al., 1999; Balu et al., 2004). The Cleland model is ideal for simulating MC behavior for a single instance of odor stimuli, and accounts for different compartments (i.e., soma, spine, dendrite). Indeed, the olfactory bulb circuit has fast dendrodendritic synaptic coupling (Rall et al., 1966) that a simple model cannot fully capture. However, large dimensional spatial models are less ideal for noise-driven systems when averaging over realizations is necessary to assess spike statistics. To this end, we have recently developed a modification of this model, removing spatial aspects and collapsing the model to a single compartment (Craft et al., 2021). Like the Cleland models, our model contains 7 ionic currents (sodium, potassium, L-type calcium, delayed rectifier, etc., see Methods) and 13 total state variables.

In this paper we study the spike statistics of our single-compartment MC model (based on Li and Cleland (2013, 2017)) with background fluctuations. Our model enables efficient yet biophysically complex simulations with which to analyze MC spiking dynamics. Our prior use of this model in Craft et al. (2021) centered around addressing how different modes of olfaction (inhale or ‘orthonasal’ versus exhale or ‘retronasal’) could manifest different operating regimes of the olfactory bulb network; we applied the model to a specific experimental data set. The general behavior of this MC model has not been investigated in detail. A natural first step is to consider the behavior of our MC model subject to background fluctuations (i.e., white noise) that is often used to mimic network and other random effects. We observe complex spiking even with constant current input, including modulation of multi-modal peaks in the interspike interval distribution **(ISI)**. With modest changes to input fluctuations (noise), the spiking variability can change dramatically even though the firing rates do not change much, exhibiting non-monotonic spike variability not commonly observed in simple spiking neuron models. We explain these dynamics with a phenomenological model of the ISI probability density function. This paper provides insights to some of the complexities of MC spiking dynamics.

## 2 Results

### 2.1 Non-standard spiking dynamics

The MC model we developed captures some real biological features exhibited in experiments. To mimic slice experiment recordings of MC, we start by analyzing the voltage traces of the MC model in response to step current injection (Fig 1). For 3 different levels of current injection (*I* = 122, 125, 130 *μ*A/cm^2^), Fig 1a clearly shows that the time to sustained spiking decreases as the current value is increased, consistent with slice experiments in Balu et al. (2004). Zoomed-in plots of the voltages (Fig 1b) shows clusters of spikes, with the number of spikes increasing with current injection value, again consistent with experiments (Balu et al., 2004). Also the time between clusters of spikes decreases with increasing current (Desmaisons et al., 1999). Our model also has subthreshold voltage values that increases with current injection value (not shown), which was reported in real MCs (Desmaisons et al., 1999). Note that our model only qualitatively captures the physiology; the described effects are more pronounced in Chen and Shepherd (1997); Desmaisons et al. (1999); Balu et al. (2004).

**Fig. 1.**
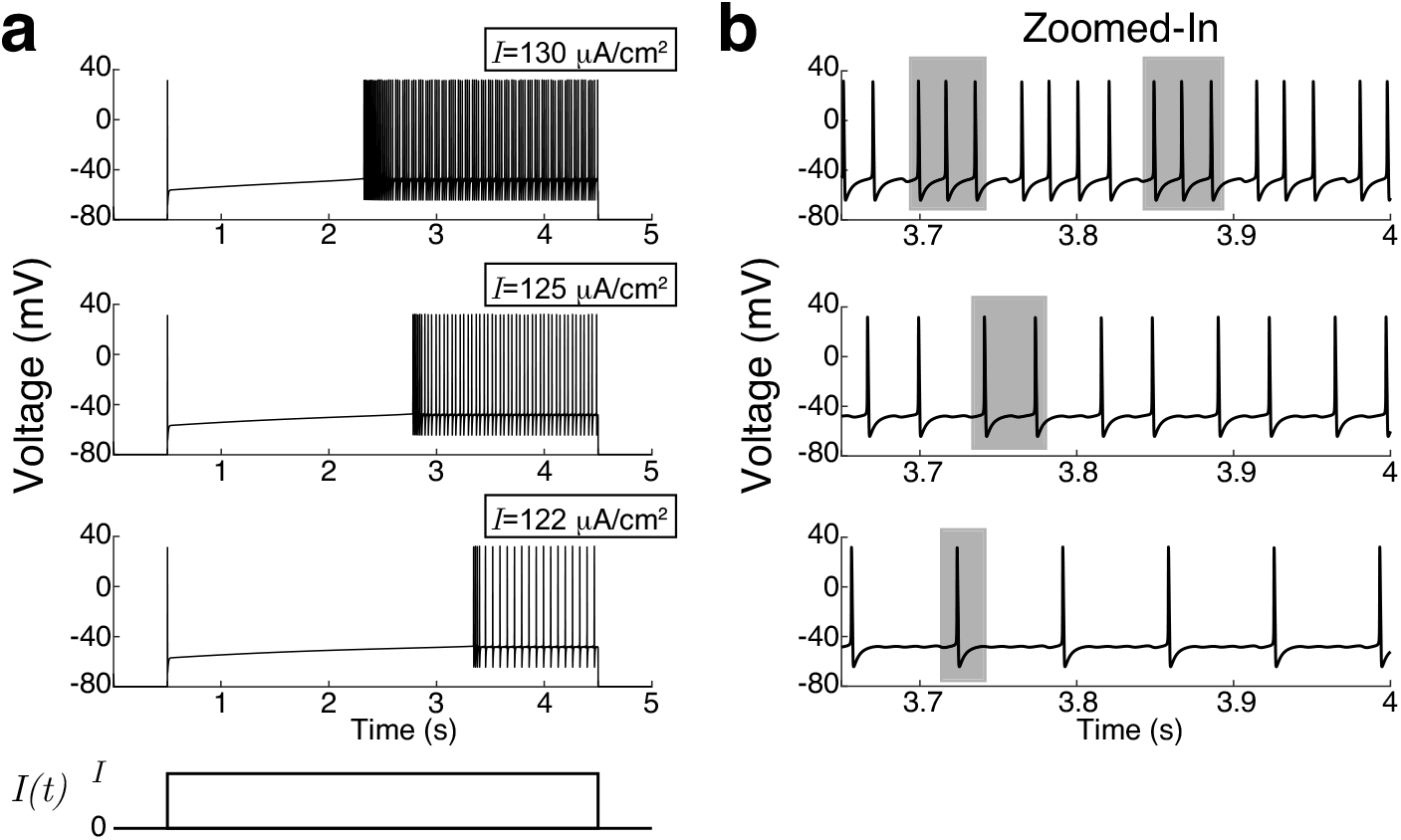
Single compartment MC model derived from a multi-compartment model captures known physiology. The transient voltage dynamics from a rest state induced by a step increase of current injection to *I*. a) The voltage traces show a time delay to spiking that increases as the current injection value I decreases, consistent with Chen and Shepherd (1997); Desmaisons et al. (1999); Balu et al. (2004) and multi-compartment model by Li and Cleland (2013). b) Zoom-in view of the voltage trajectories shows the number of spikes increases and tends to cluster with current injection value, consistent with experiments reported in Balu et al. (2004) and multi-compartment model by Li and Cleland (2013). Shaded rectangle highlights the approximate burst size. Sub-threshold oscillations are apparent (also see Fig 2a).

Another hallmark of MC voltage dynamics is sub-threshold oscillations (Chen and Shepherd, 1997; Desmaisons et al., 1999), which were not as apparent in Figure 1 because of horizontal axis scale, but evident in Figure 2a. Figure 2a shows ongoing voltage traces (long after the transient dynamics) with fixed current injection values. Notice that the temporal dynamics are complicated, with extended periods of quiescence (no spiking), rapid spiking, as well as intermittent irregular spiking (all without any background noise *σ* = 0). The multiple time-scales are certainly evident, making standard mathematical analysis of the model difficult. These complex dynamics are perhaps not surprising given that the MC model has 13 state variables.

**Fig. 2.**
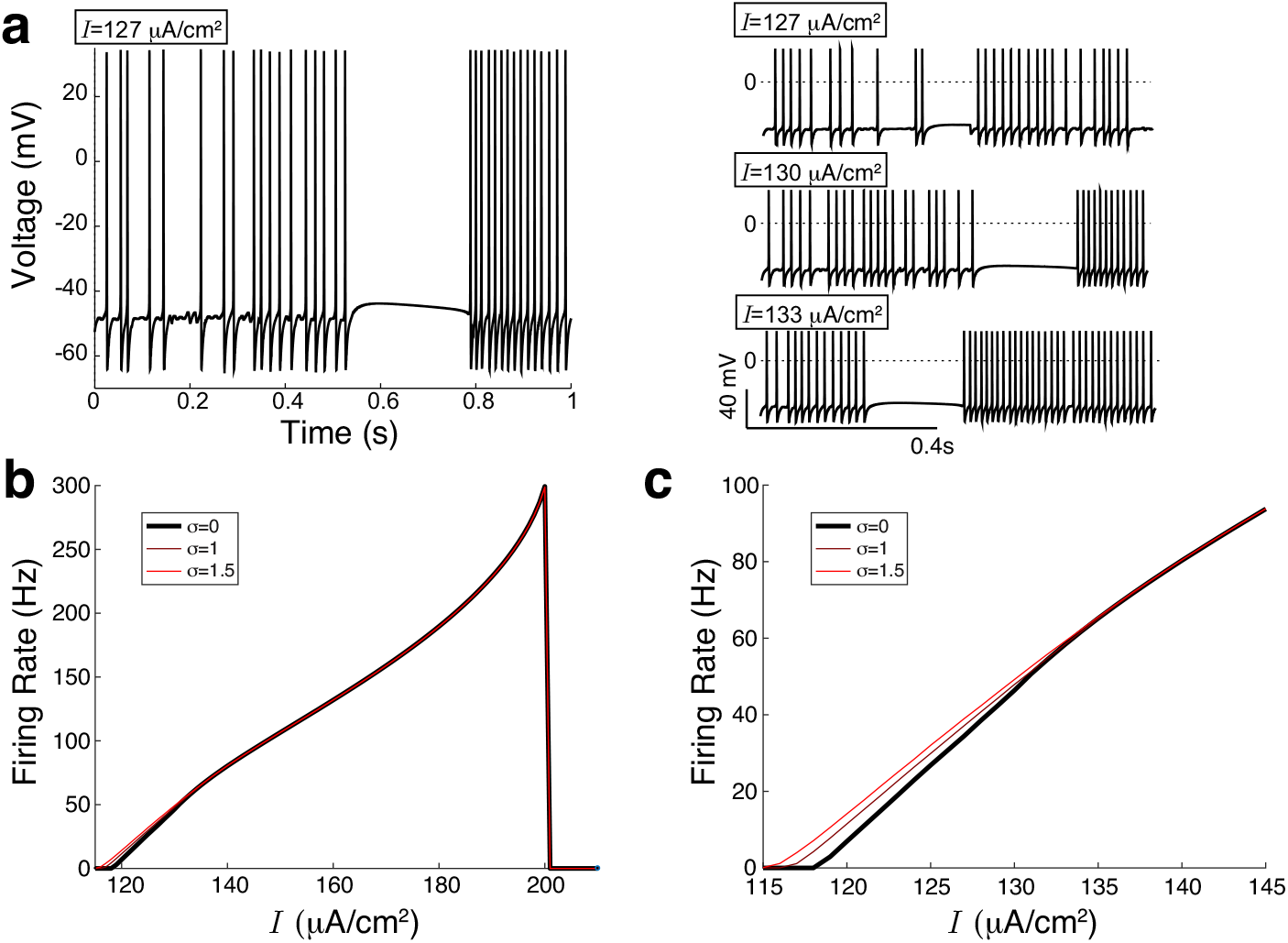
Complex spiking behavior is not well-captured by (time-averaged) firing rate. a) Ongoing voltage traces (not transient, *cf*. Fig 1) with various current values *I* and *σ* = 0. The spiking behavior has multiple time-scales with periods of quiescence, rapid and intermittent spiking. b) The FI-curve (firing rate as a function of input current) does not change much with these noise values. c) Same as b) but zoomed-in to show the slight increase in firing rate with noise.

We next consider the ‘FI-curve’ (or transfer function) of the MC model, one of the most common entities to characterize spiking behavior. This is the output firing rate (total number of spikes divided by time) as a function of input current *I*. We consider both the baseline case without noise (Fig 2b,c in black) and with relatively small noise (brown, red). Noise tends to increase the firing rates except when firing rates are very large (Lindner et al., 2003). We observe the largest changes in firing rates with increased noise where MC fires very little, consistent with many neuron models (Lindner et al., 2003). The curves in Figure 2b,c demonstrate that noise affects firing rate in a relatively simple manner, like what is observed in simpler low-dimensional spiking models, but firing rate (an average first order statistic) does not account for the aforementioned temporal dynamics in Fig 2a.

A natural entity to characterize spike statistics is the distribution of time between consecutive spikes, i.e., the ISI. The ISI has been used in many contexts (Ostojic, 2011), including classifying firing patterns from experimental data (Sacerdote et al., 2006). Multiple peaks are signatures of multiple timescales of spiking dynamics. Figure 3 shows simulations of the ISI densities for 3 fixed current values. As noise increases, it is expected for the ISI densities to shift left as well as widen (Ostojic, 2011) which is seen for *I* ∈ {120, 140} in Fig 3a & c. However, with *I* =130 *μ*A/cm^2^, the ISI has two prominent peaks that evolve into only one peak as noise increases (Fig 3b). It should be noted that the ISI for *I* = 120 *μ*A/cm^2^ when *σ* = 0 also appears to have two peaks, however, these are not as well-separated as the two peaks seen for *I* =130 *μ*A/cm^2^ when *σ* ∈ {0, 0.5}. In these plots, simulating 500,000 spikes is computationally feasible because of our single-compartment model development.

**Fig. 3.**
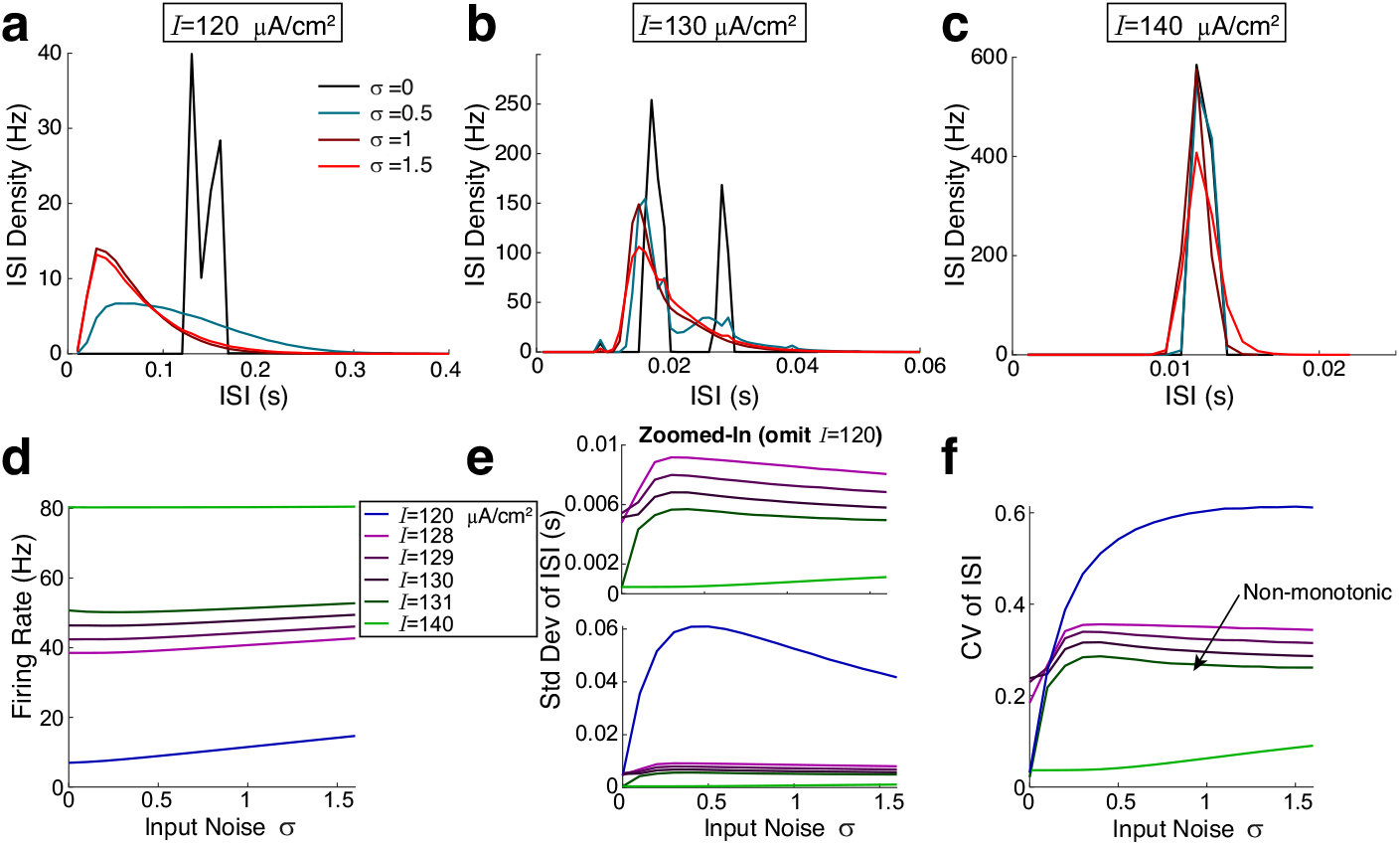
Simulation results show complex spike dynamics for physiological firing rates. a)–c) The ISI density for fixed input current and different values of input noise: *σ* = 0, 0.5,1, 1.5 using histogram bin widths of 0.01, 0.001, and 0.001s, respectively. a) (*I* = 120 *μ*A/cm^2^) and c) (*I* = 140 *μ*A/cm^2^) show expected changes with noise: the distribution shifts to the left and widens, respectively, as σ ↗ (Ostojic, 2011). b) However, with *I* = 130*μ*A/cm^2^, the ISI has 2 prominent peaks that eventually merges to 1 peak as noise increases. The ISI in a) for σ = 0 has peaks, but they are not as well-separated. d)–f) Summary ISI statistics for more σ values. d) The firing rate (inverse of mean ISI) unsurprisingly increases with noise for all *I* (although modest for *I* = 140). e) Std. dev. of ISI is a non-monotonic function of input noise σ except for large current: *I* = 140. f) The CV of ISI (std. dev. over mean) is a non-monotonic function of input noise σ for intermediate current values (*I* ∈ {128, 129, 130, 131}), coinciding with physiological firing rates.

A central focus of this paper is that the firing variability as measured by *σ_ISI_* (standard deviation of ISI) and the **C**oefficient of **V**ariation 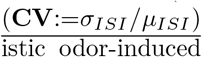 are non-monotonic functions of input noise σ for realistic odor-induced firing rates (Fig 3e,f). For intermediate current values, (*I* ∈ {128, 129, 130, 131} *μ*A/cm^2^), the firing rates are in the range of what is reported in mammals’ MC in OB *in vivo* (Cury and Uchida, 2010; Barreiro et al., 2017; Shmuel et al., 2019; Ly et al., 2021). We observe as input noise increases, the firing variability initially increases, reaches a maximum, then slowly decreases for these intermediate current values. For *I* = 120 *μ*A/cm^2^, the standard deviation of the ISI *σ_ISI_* is also non-monotonic with respect to input noise σ. However, when it is normalized by the mean via the CV, we see that it (*I* = 120) is simply a monotonically increasing function. Notice that the change in *σ_ISI_* is large, over an order of magnitude with the input noise values we used. Lastly for very high firing rates (*I* = 140 *μ*A/cm^2^), the dynamics are straight forward; uni-modal ISI with monotonically increasing spike variability with input noise.

Even with weak to modest background noise values, the multi-modal peaks in ISI density can disappear (e.g., with *I* = 130 *μ*A/cm^2^ in Fig 3b). It is thus notable that MC bursting activity with multiple time-scales (as reported in slice recordings Balu et al. (2004)) may not be operable in a network when background noise fluctuations are prominent. We do not explore the theoretical shapes of the ISIs, which may certainly have more or less modes than what is shown in Fig 3a,b,c, (a bin size was set; other bin-sizes will naturally give different shapes).

Standard neural spiking models like the leaky integrate-and-fire model do not exhibit non-monotonic spiking behavior as a function of (uncorrelated) input noise (Gerstner and Kistler, 2002). Even when the synaptic input events have temporal correlation and the spiking variability can go down with increasing input noise (see Fig 5A of Ly and Tranchina (2009)), the relationship is often monotonic – a notable exception is when there are a range of time-scales (Nesse et al., 2008).

### 2.2 Capturing results with a phenomenological model

In this section we will describe the prior observations with a simple phenomenological that will enable us to dissect how the structure of the noise and intrinsic time-scales (i.e., multi-modal ISI density) contribute to non-monotonic spiking variability.

In Appendix A, we highlight possible avenues for analysis and argue that a more formal mathematical approach to describe the observations in the previous section is likely infeasible, at least in our opinion. Although we have not discounted every possible approach to analyze our MC model, describing these results likely requires a phenomenological approach, especially considering there are 13 state variables with a range of time scales.

We approximate the ISI with the random variable *S*, governed by the equations:

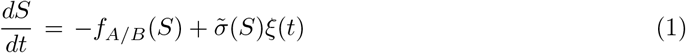

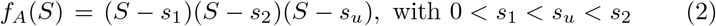

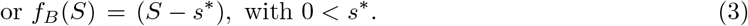

With weak noise, *f_A_* results in a multi-modal ISI with 2 peaks (Fig 3b), *f_B_* is uni-modal (Fig 3c). The advantage here is that the PDF of *S*

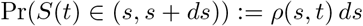

can be analytically solved for, in the steady state:

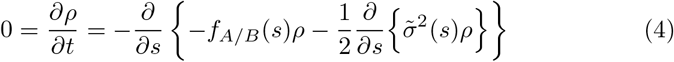

The solution is (Risken, 1989; Gardiner, 1985):

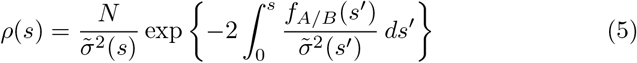

where 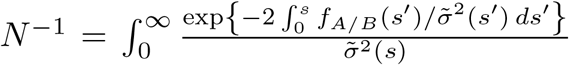 *ds* is the constant to normalize *ρ*. When 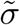 is constant, the density is simply:

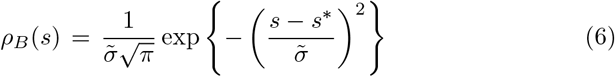

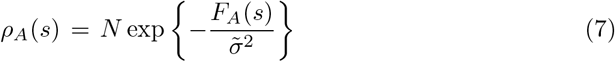

where 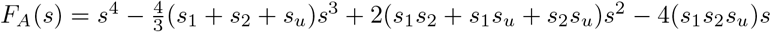 has local mins at *s*_1_, *s*_2_ (global min is(are) the point(s) furthest from *s_u_*) and a local max at *s_u_* by construction, and 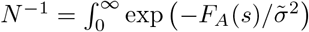 *ds* is the constant to normalize *ρ*. This formulation will thus allow us to assess how the number of peaks and the structure of the *effective* noise alters *σ_S_* and *σ_S_/μ_S_* – these are respectively the representations of σ_ISI_ and CV from the biophysical MC model.

For exposition purposes, we set the fixed points to be *s*_1_ = 4, *s*_2_ = 8, *s_u_* = 6 for *f_A_*, and *s*\* =6 for *f_B_*. First consider the case of simple additive noise where *σ* does not depend on *S*. In the uni-modal case (*f_B_*), it is not surprising that the spiking variability (*σ_ISI_, CV*) monotonically increase with 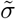 (Fig 4c,d) because the PDF (Eq (6)) is simply a Gaussian distribution with 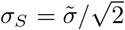 (Fig 4b). For the multi-modal case (*f_A_*), we find that for a large range of 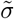, the spiking variability monotonically <u>decreases</u> with noise 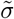 (Fig 4c,d).

**Fig. 4.**
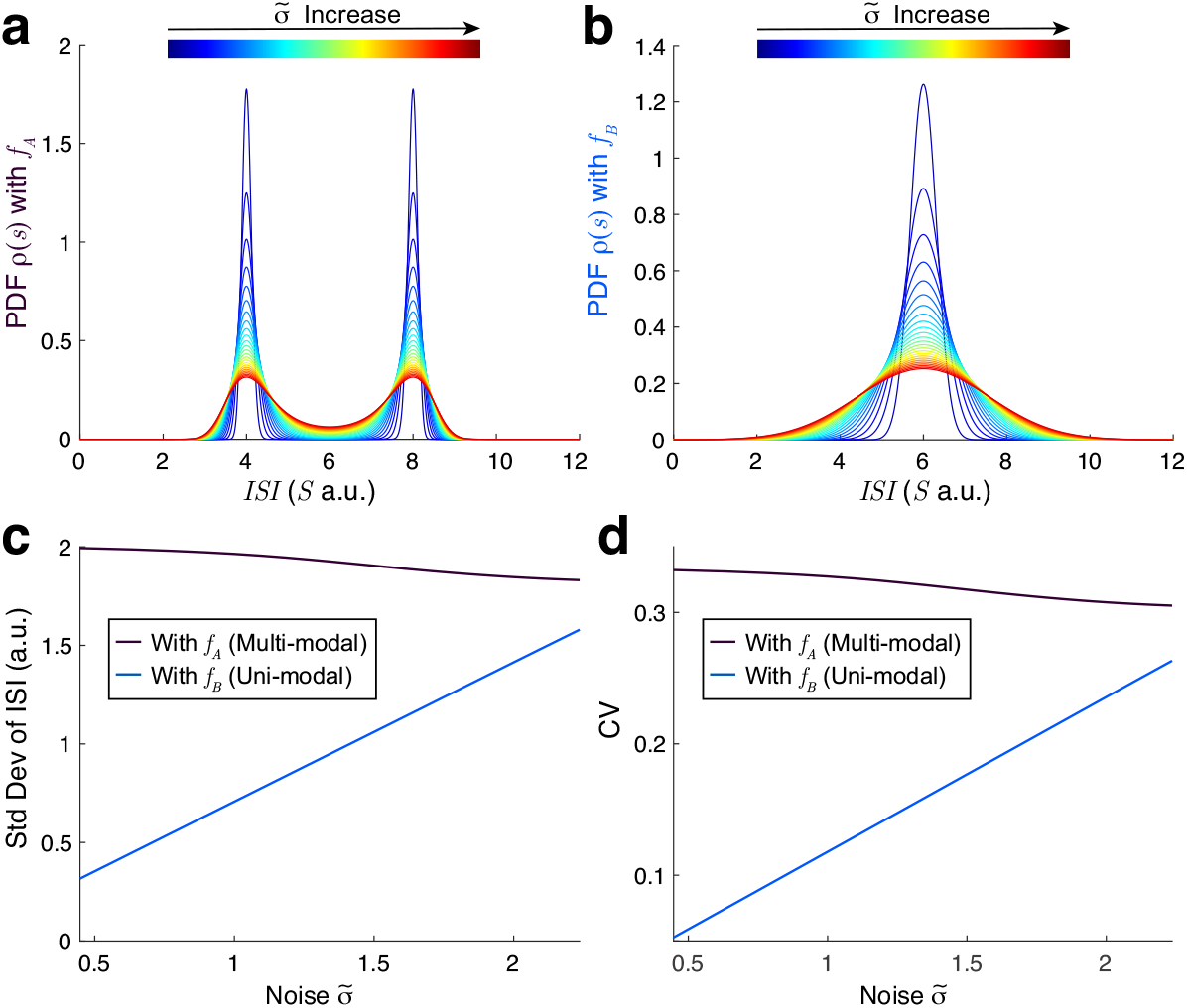
Phenomenological model of ISI (Eqs (1)–(3)) with additive noise. a) PDF of *S* with multiple stable fixed points (multi-modal), Eq (7). b) PDF of S with one stable fixed point, Eq (6). c) Resulting std. dev. of ISI (modeled with *σ_S_* and CV (d) (modeled with *σ_S_/μ_S_*). The multi-modal case (blackish) with *f_A_* shows that variability generally decreases while the uni-modal case (blueish) with *f_B_* always increases with increasing input noise.

**Fig. 5.**
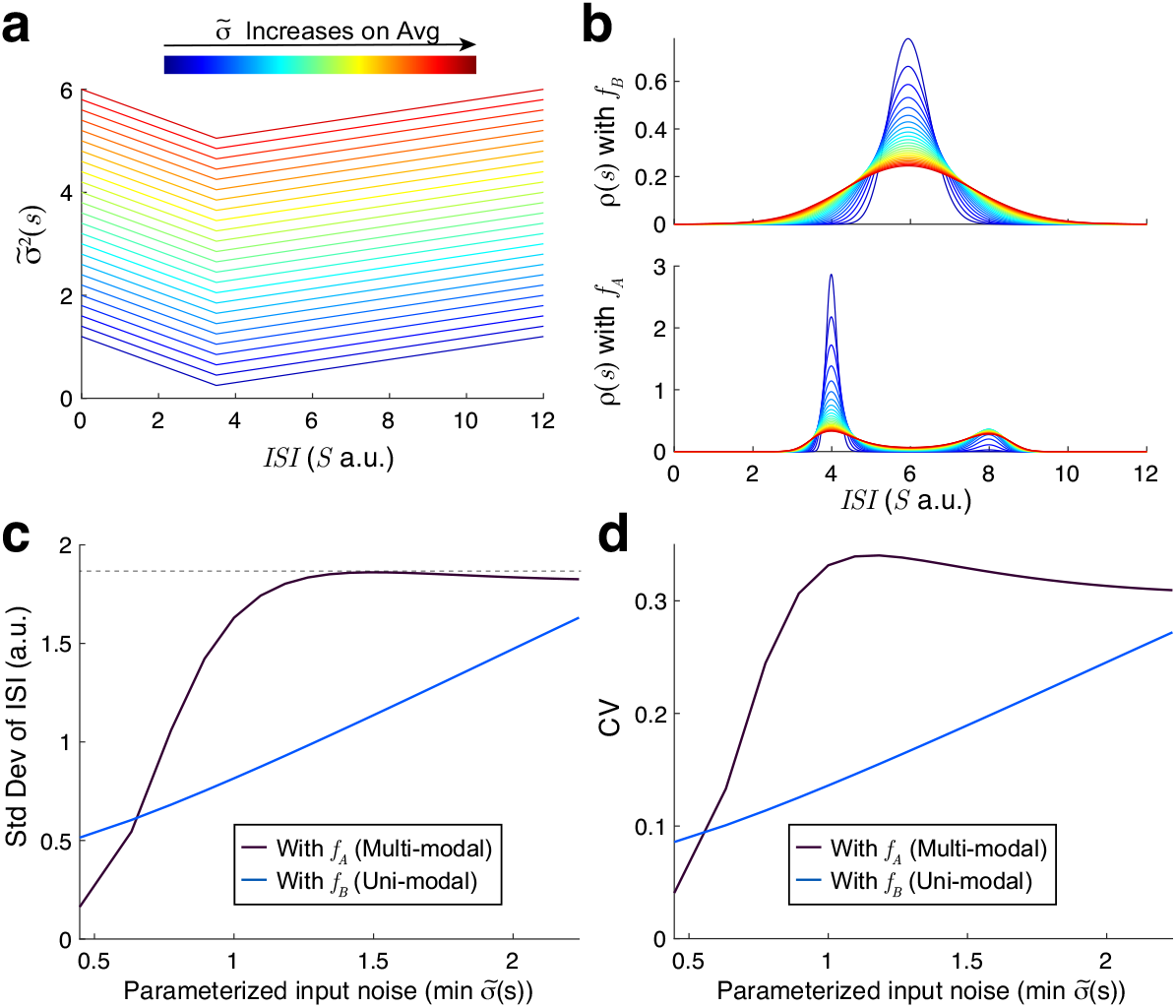
Phenomenological model of ISI (Eqs (1)–(3)) with multiplicative noise. a) Simple structure of multiplicative noise we chose (arbitrarily). b) Resulting PDFs (Eq (5)). c) The std. dev. and CV (d). The non-monotonic spiking variability is apparent with both *f_A_* (multi-modal) and multiplicative noise, while with *f_B_* (uni-modal) the resulting variability only increases.

To capture the main results in Fig 3, the results in the prior paragraph suggests that the noise 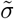 has to be multiplicative (i.e., depend on *S*) in this specific model. Since increasing input noise generally results in a gradual diminishing of multi-modal peaks to uni-modal, we set 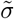 in Eq (5) to be an inverted triangle with minimum at *s* = 3, and model increase in noise by an additive shift (see Fig 5a). These choices are arbitrary, but they nonetheless capture the changes in the ISI distribution (Fig 5c,d) observed in the biophysical model (Fig 3e,f; the right peak broadens and flattens while left peak’s height decreases as input noise increases).

Importantly, the uni-modal case (*f_B_*) generally results in increases in spiking variability with noise, while the multi-modal case (*f_A_*) exhibits an increase followed by a decrease in spiking variability as noise increase.

Taken together, we conclude that both the multi-modal peaks in the ISI and the (effective) multiplicative input noise (or ISI dependent) noise are crucial for the results we observe in this specific model. However, it is important to note that this observation is not a general fact; i.e., not all models with uni-modal ISI densities behave the same way as this scalar phenomenological model with input noise. Specifically, Nesse et al. (2008) used a simple Fitzhugh-Nagumo model with adaptation endowed with a large range of time-scales and found non-monotonic CV of spiking as input noise (and applied current) increased, with uni-modal ISI densities. Their work identifies the range of time-scales as a key signature in non-monotonic spiking CV, because when their adaptation variable has only one time-scale, the applied current (and perhaps input noise) is monotonically related to CV. See Discussion below and Appendix A for further context.

We close with remarks about the biophysical connection and the generality of the phenomenological model results. In our specific model, the multiplicative noise 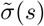 does not have a direct connection to the biophysical MC model. But note that it is very common to have multiplicative noise in a reduced/- transformed model from a larger biophysical model that originally has additive noise in the voltage. For example, see equation for phase reduced model (Eq (A1)). We decided to use a state dependent 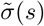 for simplicity that did not require wholesale changes to the model above (Eq (1)–(3)). In our simple model (Eq (1)–(3)) the effect of 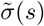 is to broaden and flatten the right peak while lowering the height of the left peak while fixing the rest of the parameters. However, note that other scalar models with <u>additive noise</u> can capture the non-monotonic Std. Dev. and CV (increase followed by a decrease). For example, consider a mixture of two Gaussians with five parameters:

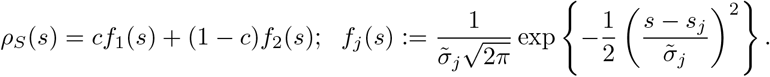

After extensive numerical investigations, we found the simplest way to capture non-monotonic Std. Dev. and CV (increase followed by a decrease) is via: with the same *s_j_*, let 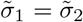 be the same additive input noise as in Fig 4a,b, and let c decrease: 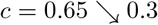.

## 3 Discussion

This paper provides insights to MC spiking variability modulation that relies first on a single-compartment reduction to pragmatically simulate large numbers of spikes, and then on a phenomenological framework to assess how input noise alters the output spike interval distribution. We find that the temporal dynamics of spiking can significantly change with input noise and that the spiking variability (measured by ISI) does not monotonically increase with biophysically realistic input noise level. These dynamics may impact odor processing as MCs are an important type of relay cells in the olfactory bulb that propagate odor information to downstream cortical brain regions. We remark that tufted cells, which we did not explicitly consider here, are also known to have a similar function. There is a recently developed tufted cell model (Viertel and Borisyuk, 2019) that may be simple enough to analyze in the same vein as here, and might be considered in future research.

Although we do not consider a comparison of an odor processing task between: i) MC model with a monotonically increasing relationship between spike variability and input noise, and ii) MC model with a non-monotonic relationship, we highlight possible implications of our results on OB odor processing. The input noise to MC comes from numerous sources, including olfactory receptor neurons (**ORNs**) from upstream, and within OB there are granule, periglomerular cells, and excitatory interneurons that are presynaptic to MC. So how the input noise precisely varies for a specific odor task is difficult to determine, but it is plausible that the input noise increases with odor compared to spontaneous activity via ORN inputs (Friedrich and Laurent, 2004) – input noise could also change when there are multiple similar odors compared to a single odor (Rospars et al., 2008), and in a multitude of ways. The MC spiking variability across trials is important for olfactory processing – awake behaving rats show reliable MC spiking for a given odor (Cury and Uchida, 2010) with relatively small trial variability (see Fig 1D, 2A, 2C in Cury and Uchida (2010)). In zebrafish, Friedrich and Laurent (2004) showed that MC spike variability (across trials) increased after odor presentation followed by dramatic decreases, which coincided with better odor identification as time proceeded from odor presentation. We did not consider populations of MC, but Friedrich and Laurent (2004) found that decorrelation across populations of MCs played a large role in odor identification, consistent with Wanner and Friedrich (2020). Therefore, how MC spike variability modulates as input noise increases, particularly when a decrease in MC spiking variability is possible with increased input noise for physiologically relevant firing rates, could be important for odor processing.

### 3.1 Analyzing simulation results

There is often a trade-off between mathematical analysis and biological realism in model analysis, our results here focus on a realistic MC model. Some modelers have used simple MC models ((Zhou et al., 2013; Marella and Ermentrout, 2008, 2010) and even very recently: (Barreiro et al., 2017; Pyzza et al., 2021; Patel and Rangan, 2021)) that, despite the virtue of these works, could result in overlooking important MC spiking behavior when the spike variability is crucial.

Describing the complexities of the ISI density in our single cell MC model with a quantitatively accurate reduction method is elusive, at least to the best of our knowledge. With constant current input, phase reductions methods are commonly used to analytically capture spiking dynamics when the time between spikes are relatively regular (which does not hold in the models here). Using such approaches to approximate the dynamics with a single scalar variable would yield approximations where the spiking variability monotonically increases with input noise (Ly and Ermentrout, 2011) (see Appendix A), in contrast to our results. There have recently been a plethora of advances in phase reduction theory using more than one state variable (i.e., amplitude variables, not just phase) that can sometimes be viable for analysis (Wilson and Ermentrout, 2019; Ashwin et al., 2016; Wedgwood et al., 2013). For example, the standard assumptions of weak coupling (Wilson and Ermentrout, 2019) and noise (Thomas and Lindner, 2014; Schwabedal and Pikovsky, 2013) can be relaxed in some of these formulations. However, the assumption of (nearly) periodic dynamics must still hold in all of the methods mentioned. These contemporary phase reduction methods generally require two or more state variables which often rely on numerical simulations anyway – we are unaware of any phase reduction theory where the approximate ISI density is readily available.

Another common analytic approach to understanding the effects of weak noise on (output) spiking in excitable systems is using the Arrhenius escape rate, i.e., potential well models, that have been successfully applied to many systems (Nesse et al., 2008; Ly and Doiron, 2017; Plesser and Gerstner, 2000; DeVille et al., 2006; Lundstrom et al., 2009; Kramers, 1940). Many of the derived formulas for the ISI distribution (see Appendix A for more details) in these prior works are too simplistic for our MC model. However, using a separation of time-scales, Nesse et al. (2008) were able to analytically describe, with an Arrhenius-escape framework, the spiking dynamics of an excitable Fitzhugh-Nagumo model with multiple time-scales in the adaptation variable. Indeed, Nesse et al. (2008) found a non-monotonic relationship of the CV with input noise (in their Fig 2, see cross-section with a fixed *I*). Their approach applied to our MC model would require identifying all of the effective timescales in our 13 variable model *and* having a significant separation of timescales, then the slower variables are assumed to be fixed (then slowly varying) and the faster variables are included in the potential dynamics (see Ly and Doiron (2017) too, also see Appendix A for more details). The viability and the accuracy of this approach for capturing our results is an open question but beyond the scope of this current study. Our description of this particular nonmonotonic relationship of escape rate (or spiking) variability with input noise in a large dimensional model compliments the results of Nesse et al. (2008) as an understudied phenomena not observed in standard/simple spiking models.

The result that the spiking variability is non-monotonic and maximal at an intermediate input noise value (for physiological firing rates) might seem related to the stochastic resonance phenomena that has been observed and analyzed in many areas of science (Gammaitoni et al., 1998). Stochastic resonance refers to an optimal (maximum) response (i.e., firing or signal to noise) at an intermediate noise level (Lindner, 2002); or the related coherence resonance where there is an optimal frequency of sinusoidal input to illicit maximum responses (Reinker et al., 2003, 2006; Lindner, 2002). Firing rate response is different than spiking variability as measured by *σ_ISI_* and CV. Although note that Lindner (2002) showed an optimal level of input noise could minimize CV in a Fitzhugh-Nagumo model, which is the opposite of what we observe here (also see Appendix A).

Although we have observed a non-monotonic relationship between simple uncorrelated white noise input and output spiking variability (i.e., CV), note that there are other studies that have described dependence of CV on more complicated input types (e.g., with temporal correlation or adaptation and other features (Schwalger and Schimansky-Geier, 2008; Schwabedal and Pikovsky, 2013)). We have considered a specific biological model for an important olfactory bulb cell, in contrast to these other papers that have analyzed general spiking models with some attributes – whether they are directly related to this paper is unlikely in our opinion, but could be explored in future research. Beyond the ISI, there are other statistical measures of a spike train, such as the autocorrelation and power spectrum, that could be unrelated to the ISI histogram (see Appendix B).

### 3.2 Limitations

There are several limitations of the work here. First, in the actual neural network, MCs are known to be coupled to other specific cells in the bulb (e.g., granule and periglomerular cells and other excitatory interneurons), and receive feedback input from piriform cortex. Related to this point, stochastic inputs can at times have more complex structure than white noise, even when considering an average over many cells. The synaptic coupling in the olfactory bulb is distinct from other neural networks, mediated by dendrodendritic interactions that results in faster coupling than the standard axon-to-dendrite coupling. These and other physiological details could change our results, at least quantitatively. Considering such extensions is an interesting avenue for future work.

## 4 Material and Methods

See GitHub page https://github.com/michellecraft64/MCuncoup for freely available code simulating the models in this paper.

Our MC model is based on a multi-compartment model developed by the Cleland Lab Li and Cleland (2013, 2017), where all compartments (soma, dendrite, axons, spin) are combined into a single compartment, as in Craft et al. (2021). All parameters and function for intrinsic ionic currents and their gating variables are the same as in Li and Cleland (2013, 2017) except the maximal conductance, which is set to be the sum of all maximal conductance values from all compartments. Excluding the many auxiliary functions, there are a total of 13 state variables (ODEs).

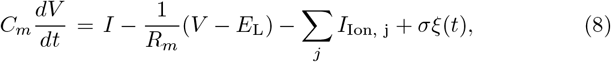

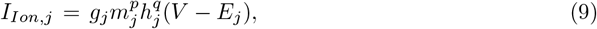

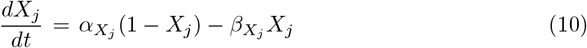

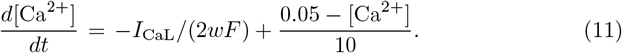

The variable *X* (Eq (10)) generically represents 11 different gating variables for the 7 different ionic currents (see Table 1), and the transition rates *α, β* can depend on *V*, [Ca^2+^], etc. (see Table 2 for gating variable equations). Note, *I*_NaP_ current is not represented by Eq (10) since it is defined by *m* = *m*_∞_(*V*). The noise level is measured by *σ* ≥ 0, and *ξ*(*t*) is white noise with 〈*ξ*(*t*)〉 = 0 and 〈*ξ*(*t*)*ξ*(*t*′)〉 = *δ*(*t* – *t*′).

**Table 1.** Description of model parameters and values. Each of these values are the same as defined by Li and Cleland (2013, 2017); see Craft et al. (2021) for the same reduction. One minor difference with Craft et al. (2021) is that *gDR* = 15mS/cm^2^ here, but was set to *gDR* = 75mS/cm^2^ in Craft et al. (2021).

| Resistance and Capacitance |  |  |
| --- | --- | --- |
| Description | Variable | Value |
| Membrane Resistance | $R_m$ | $30 \text{ k}\Omega/\text{cm}^2$ |
| Membrane Capacitance | $C_m$ | $1.2 \text{ }\mu\text{F}/\text{cm}^2$ |
| Ionic Currents |  |  |
| Description | Variable | Max Conduct. Value ($\text{mS}/\text{cm}^2$) |
| Fast, Spike-Generating Sodium Current | $I_{Na}$ | $g_{Na} = 120$ |
| Persistent Sodium Current | $I_{NaP}$ | $g_{NaP} = 0.42$ |
| Potassium Delayed Rectifier | $I_{DR}$ | $g_{DR} = 15$ |
| Fast-Activating Transient Potassium Current | $I_A$ | $g_A = 10$ |
| Slow-Inactivating Transient Potassium Current | $I_{KS}$ | $g_{KS} = 84$ |
| L-type Calcium Current | $I_{CaL}$ | $g_{CaL} = 0.85$ |
| $\text{Ca}^{2+}$-Dependent Potassium Current | $I_{KCa}$ | $g_{KCa} = 5$ |
| Reversal Potentials |  |  |
| Description | Variable | Value (mV) |
| Leak Current Reversal Potential | $E_L$ | -60 |
| Sodium Reversal Potential | $E_{Na}$ | 45 |
| Potassium Reversal Potential | $E_K$ | -80 |
| Calcium Reversal Potential | $E_{Ca}$ | $\frac{RT}{2F} \log \left( \frac{10}{[\text{Ca}^{2+}]} \right)$ |
| Calcium Dynamics |  |  |
| Description | Variable | Value |
| Perimembrane Thickness | $w$ | 1 micron |
| Boltzman Constant * Temperature / Faraday Constant | $\frac{RT}{F}$ | 26.55 mV |
| $\text{Ca}^{2+}$ Removal Rate | $\tau_{Ca}$ | 10 ms |
| Intracellular $\text{Ca}^{2+}$ Concentration | $[\text{Ca}^{2+}]$ | dynamic |
| $\text{Ca}^{2+}$ Resting Concentration | $[\text{Ca}^{2+}]_{\text{rest}}$ | $0.05 \text{ }\mu\text{mol/l}$ |

**Table 2.** Gating variable dynamics. Same dynamics as was used in Li and Cleland (2013), except we used a function fit for *I_DR_* rather than a lookup table. When gating variables are given as *X*_∞_ and *τ_X_*, the equation is: 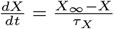 rather than (10). The last column refers to the following references: [1] Bhalla and Bower (1993), [2] Migliore et al. (2005), [3] Wang (1993).

| Ionic Current | Gating Var | $\alpha_x$ or $x_\infty$ | $\beta_x$ or $\tau_x$ (ms) | Source |
| --- | --- | --- | --- | --- |
| $I_{Na}$ | $p = 3$<br>$q = 1$ | $\alpha_m = \frac{0.32(V+45)}{1 - \exp(-(V+45)/4)}$<br>$\alpha_h = \frac{.128}{\exp((V+41)/18)}$ | $\beta_m = \frac{-.28(V+18)}{1 - \exp((V+18)/5)}$<br>$\beta_h = \frac{4}{1 + \exp(-(V+18)/5)}$ | 1 |
| $I_{NaP}$ | $p = 1$ | $m_\infty = \frac{1}{1 + \exp(-(V+50)/5)}$ | | 3 |
| $I_{DR}$ | $p = 2$<br>$q = 1$ | $m_\infty = \frac{((V+100)/150)^{8.585}}{.575^{8.585} + ((V+100)/150)^{8.585}}$<br>$h_\infty = .433(1 + \tanh(-\frac{V+13.925}{13.02})) + .1337$ | $\tau_m = \frac{\exp(-(V+30)/66.378)}{.27654} + \frac{2.89}{1 + \exp(-(V-19.0524)/12.879)}$<br>$\tau_h = 50$ | 1 |
| $I_A$ | $p = 1$<br>$q = 1$ | $m_\infty = \frac{1}{1 + \exp(-(V-17.5)/14)}$<br>$h_\infty = \frac{1}{1 + \exp((V+41.7)/6)}$ | $\tau_m = \frac{25 \exp((V+45)/13.3)}{3.3(1 + \exp((V+45)/10))}$<br>$\tau_h = \frac{55.5 \exp((V+70)/5.1)}{3.3(1 + \exp((V+70)/5))}$ | 2 |
| $I_{KS}$ | $p = 1$<br>$q = 1$ | $m_\infty = \frac{1}{1 + \exp(-(V+34)/6.5)}$<br>$h_\infty = \frac{1}{1 + \exp((V+68)/6.6)}$ | $\tau_m = 10$<br>$\tau_h = 200 + \frac{330}{1 + \exp(-(V+71.6)/6.85)}$ | 3 |
| $I_{CaL}$ | $p = 1$<br>$q = 1$ | $\alpha_m = \frac{7.5}{1 + \exp(-(V-13)/17)}$<br>$\alpha_h = \frac{6.8 \times 10^{-3}}{1 + \exp((V+30)/12)}$ | $\beta_m = \frac{1.65}{1 + \exp(-(V-14)/4)}$<br>$\beta_h = \frac{.06}{1 + \exp(-(V/11))}$ | 1 |
| $I_{KCa}$ | $p = 1$ | $\alpha_m = \frac{-500 \exp((V-65)/27)(0.015 - [\text{Ca}^{2+}])}{1 - \exp(-([\text{Ca}^{2+}] - .015)/.0013)}$ | $\beta_m = 0.05$ | 1 |

## Declarations

## Acknowledgments

We thank Woodrow Shew, Shree Hari Gautam, and Andrea Barreiro for many conversations about olfactory bulb cells and circuits.

## Code availability

The code to implement the models is freely available at https://github.com/michellecraft64/MCuncoup.

## Data availability

This study did not use any experimental data.

## Author contributions

All authors contributed to the study conception and design. Material preparation and analysis were performed by Cheng Ly, and Michelle Craft. The first draft of the manuscript was written by Cheng Ly and Michelle Craft and all authors commented on previous versions of the manuscript. All authors read and approved the final manuscript.

## Appendix A Viability of alternative approaches

A common approach to reduce the number of state variables in describing (regular) spiking is to apply a phase reduction, which seems appealing because the results hold for relatively weak noise forcing where firing rates do not vary much. This motivated us to analyze the bifurcation between quiescence and spiking (using XPP-AUTO (Ermentrout, 2002; Doedel, 1981)), in the noiseless case (Fig A1a). We see that the stable rest state loses stability via a saddle-node on invariant circle (**SNIC**) bifurcation as the applied current increases. However, we do not find actual periodic solutions for relevant firing rates with which to calculate commonly used entities for analysis like the Phase-Resetting Curve (**PRC**). Curiously, the bifurcation diagram shows stable period solutions (green dots) that are interlaced with unstable periodic solutions. Figure A1b shows that even with *I*_app_ = 144 *μ*A/cm^2^ where the diagram (Fig A1a) suggests there is a well-behaved periodic solution, the voltage (and other variables) are not strictly periodic. Note that with *I* = 144, the firing rates are large (91.13Hz) and not physiologically relevant. Also, for *I* = 144 although the PRC is numerically calculable (Fig A1c), the rather large negative region does not resemble canonical PRCs associated with a SNIC (Rinzel and Ermentrout, 1989; Ermentrout, 1996).

Note that in a standard phase reduced scalar model (Ermentrout and Terman, 2010):

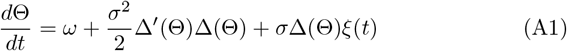

the ISI density can be approximated (assuming weak noise) via (Ly and Ermentrout, 2011):

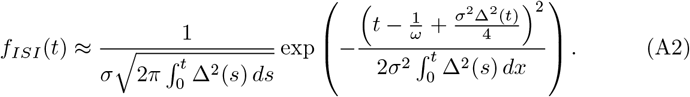

**Fig. A1.**
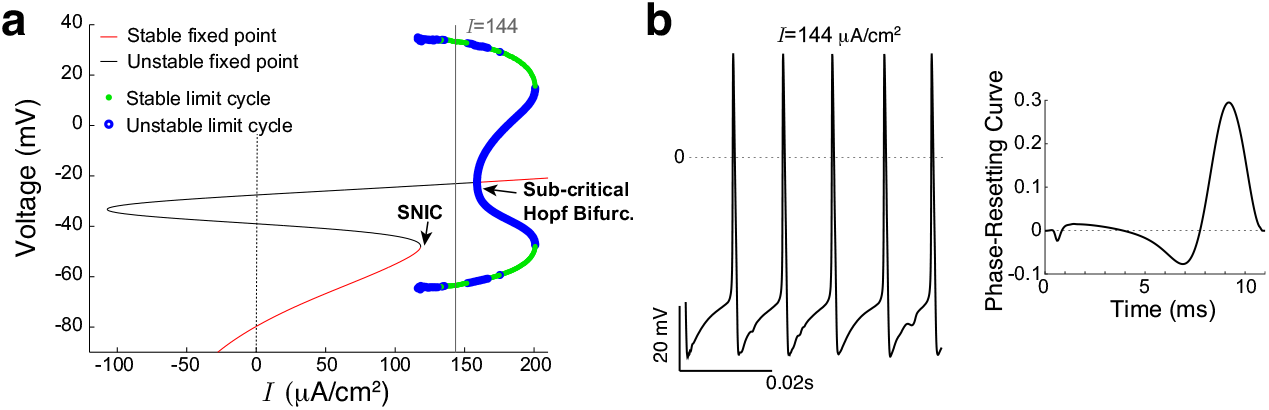
Dynamics of noiseless MC model: phase reduction assumptions violated. a) Bifurcation diagram of voltage varying *I* shows SNIC at onset of spiking. b) The voltage traces in Fig2a are not strictly periodic, even for an ideal well-behaved *I* value. For unrealistically high firing rates (*I* = 144 *μ*A/cm^2^, firing rate: 91.13Hz), we see the system is not periodic despite the bifurcation diagram suggesting it should be. The infinitesimal Phase-resetting curve can be numerically calculated with XPP (Ermentrout, 2002); notice that the negative region is rather large, not resembling canonical PRCs synonymous with a SNIC (Rinzel and Ermentrout, 1989; Ermentrout, 1996). This all suggests phase reduction descriptions would likely be inadequate to capture the observed phenomena in the regimes we are interested in (i.e., smaller *I* with physiological firing rates).

Since the formula closely resembles a normal distribution, the *σ_ISI_* generally increases as 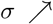. Even though there is multiplicative noise: *σ*Δ(Θ), these models would not help describe observations in Figure 3.

Another common approach to model analysis with weak noise is use a potential well: 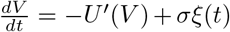, where *U* is the potential function, either principally derived from the system (simple) or ad-hoc (high-dimensional). E.g., for the leaky-integrate-and-fire model, 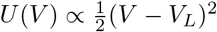 and *V* has a stable fixed point at the global minimum *V* = *V_L_*. This approach was pioneered in physics by Kramers (1940) and has been applied to several neural models where “exiting” from the potential well from crossing a threshold is spiking. The rate of spiking is often ∝ *e*^-*U/σ*2^, which is not directly related to spiking variability. The signal–to–noise ratio (**SNR**) in these systems, and in other applications of stochastic resonance, often have a maximal SNR value for an intermediate level of noise (Gammaitoni et al., 1998). However, this dynamic is associated with a minimum variability value in the denominator (ignoring the dynamics of the signal in the numerator) of SNR, rather than a maximal spiking variability for intermediate input noise level, as we have observed in the MC model.

Whether the potential well or “Arrhenius escape” approach by Nesse et al. (2008) for a low-dimensional Fitzhugh-Nagumo model (with 1 activity variable x and a set of identical adaptation variables *H* endowed with multiple timescales that depend on *x*) could be successfully applied to our MC model is an open question. Nesse et al. (2008) exploited a separation of time scales, the slow variable was fixed and the mean first passage time (or escape) *T* could be calculated (in the fast variable) and set to the inverse of the mean firing rate: λ(*H*(*t*)) = 1/*T*(*H*(*t*)). The ISI density is approximated with:

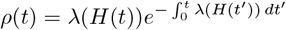

we see how the slow variation in *H* affects *ρ*(*t*). This framework successfully described the non-monotonic spiking dynamics (in the CV at least) in their model. As previously mentioned in the Discussion, an analogous approach would require identifying all of the effective time-scales in our 13 variable model *and* having a significant separation of time-scales when the neuron is excitable. Even if the slow variables are frozen, one would still have to calculate the mean first passage time with the remaining fast variables, which is generally not feasible unless the resulting dimension is small. Solving for the mean first passage time requires solving an ODE system derived from the backward Fokker-Planck equation, a PDE with the number of dimensions equal to the number of state variables (Gardiner, 1985; Risken, 1989). The viability and the accuracy of this approach for capturing our results is an open question but beyond the scope of this current study.

## Appendix B Other spike statistics

We have focused on the ISI distribution, but there are other commonly used entities to characterize neural spike trains. For instance, the autocorrelation function (*ACF*) and power spectrum (*P*), defined below, are commonly used and can be unrelated to the ISI, in particular when the system does not reset after a spike. Letting *R*(*t*) denote the spike train consisting of 0’s and 1’s, the (normalized) autocorrelation function is:

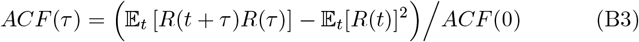

and the power spectrum:

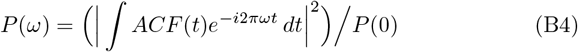

Figure B2 shows these entities for the biophysical MC model with various applied current and input noise values.

With *I* = 120*μ*A/cm^2^, *ACF*(*τ*) is relatively flat for these input noise values, while *P*(*ω*) changes from having peaks at regularly spaced intervals with no noise (black) to being relatively flat with 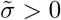. With *I* = 130 *μ*A/cm^2^, *ACF*(*τ*) has local maximas at irregularly spaced *τ* with no noise (black) that flatten out as input noise increases; the *P*(*ω*) is similar to *I* = 120, *μ*A/cm^2^ but the curves have smaller values compared to *I* = 120. With *I* = 140 *μ*A/cm^2^, *ACF*(*τ*) indicates relatively regular spiking, although increased input noise shifts and broadens the peaks (same for *P*(*ω*)). The flatter *ACF* with *I* = 120 compared to larger *I* indicates that the spiking has less temporal regularity, which is not surprising. For a given value of *I* (i.e., a row in Fig B2), increasing input noise flattens the *ACF*(*τ*) and shifts and/or diminishes peaks (local max), and for *P*(*ω*) input noise can broaden/shift/diminish peaks. Overall, the effects of input noise are nonlinear and highly dependent on *I*.

**Fig. B2.**
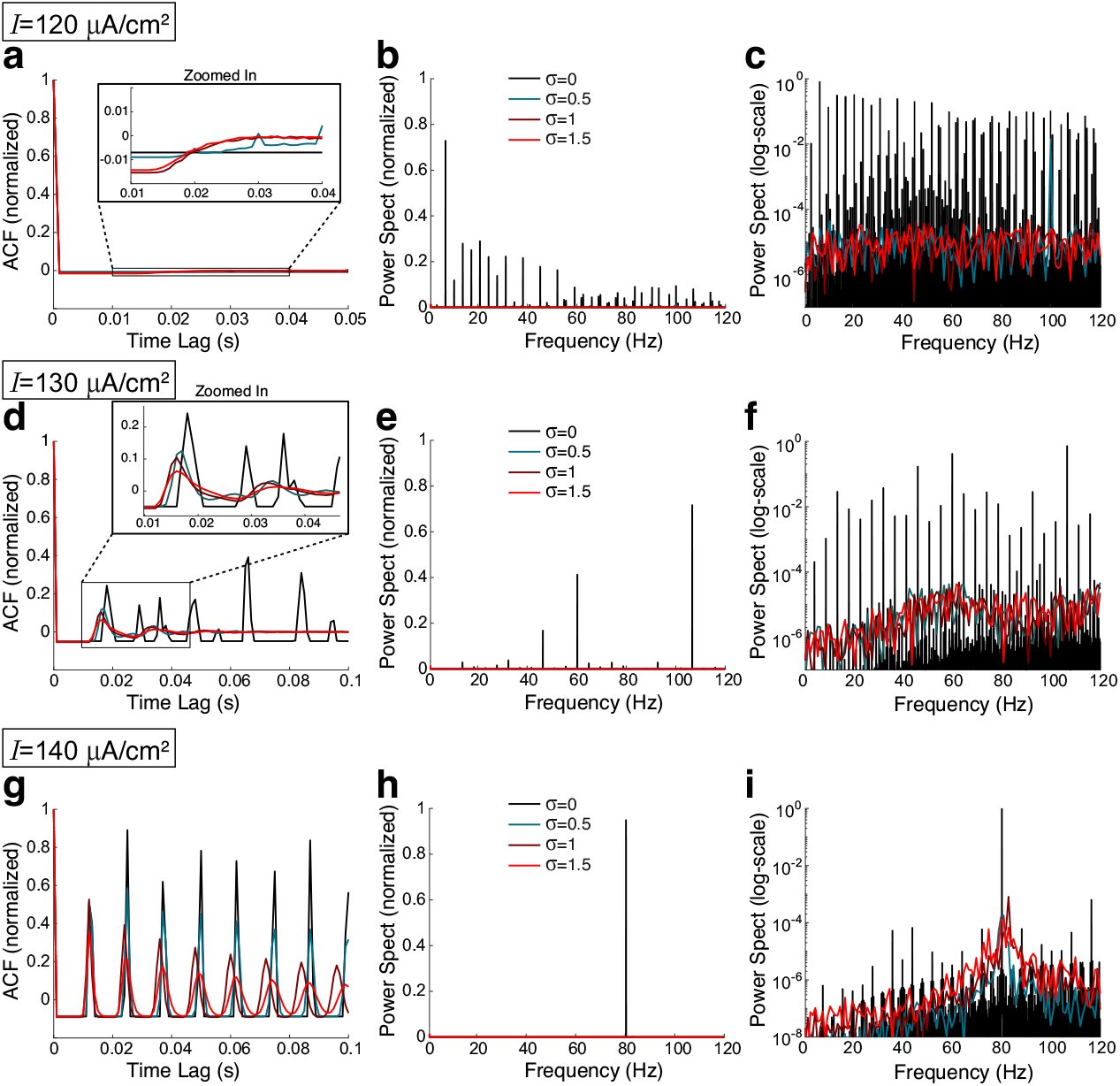
The autocorrelation function (Eq (B3)) and power spectrum (Eq (B4)) of the MC model. Three values of input current: *I* = 120*μ*A/cm^2^ in a)–c), *I* = 130*μ*A/cm^2^ in d)-f), *I* = 140 *μ*A/cm^2^ in g)-i), with each panel showing the effects of increasing input noise σ. The effects of input noise are nonlinear and highly dependent on *I*.

The *ACF* and *P* were plotted using built-in functions in MATLAB, and there appears to be slight numerical round-off errors in the *ACF*, e.g., between the peaks in Fig B2g, *ACF* ≈ 0, and likely in the *P*. Nevertheless, these plots given insight to some dynamics of the various spike trains.

